# Genomic and phenotypic evidence for an incomplete domestication of South American grain amaranth (*Amaranthus caudatus*)

**DOI:** 10.1101/025866

**Authors:** Markus G. Stetter, Thomas Müller, Karl J. Schmid

## Abstract

The process of domestication leads to major morphological and genetic changes, which in combination are known as domestication syndrome that differentiates crops from their wild ancestors. We characterized the genomic and phenotypic diversity of the South American grain amaranth *Amaranthus caudatus*, which has been cultivated for thousands of years and is one of the three native grain amaranths of South and Central America. Previously, several models of domestication were proposed including a domestication from the close relatives and putative ancestors *A. hybridus* or *A. quitensis*. To investigate the evolutionary relationship of *A. caudatus* and its two close relatives, we genotyped 119 amaranth accessions of the three species from the Andean region using genotyping-by-sequencing (GBS) and compared phenotypic variation in two domestication-related traits, seed size and seed color. The analysis of 9,485 SNPs revealed a strong genetic differentiation of cultivated *A. caudatus* from the relatives *A. hybridus* and *A. quitensis*. The two relatives did not cluster according to the species assigment but formed mixed groups according to their geographic origin in Ecuador and Peru, respectively. *A. caudatus* had a higher genetic diversity than its close relatives and shared a high proportion of polymorphisms with their wild relatives consistent with the absence of a strong bottleneck or a high level of recent gene flow. Genome sizes and seed sizes were not significantly different between *A. caudatus* and its relatives, although a genetically distinct group of *A. caudatus* from Bolivia had significantly larger seeds. We conclude that despite a long history of human cultivation and selection for white grain color, *A. caudatus* shows a weak genomic and phenotypic domestication syndrome and is an incompletely domesticated species because of weak selection or high levels of gene flow from its sympatric close undomesticated relatives that counteracted the fixation of key domestication traits.

## Introduction

Research on the domestication of crop plants revealed that numerous traits can be affected by domestication, which results in so-called domestication syndromes. The type and extent of domestication syndromes depends on the life history and uses of crop plants (Meyer *et al* 2012), although crops from distantly related plant families frequently show similar domestication phenotypes. For example, the ‘classical’ domestication syndrome, which includes larger seeds, loss of seed shattering, reduced branching, loss of seed dormancy and decreased photoperiod sensitivity, is observed in legumes and grasses (Abbo *et al* 2014; Hake & Ross-Ibarra, 2015). Similar to phenotypic diversity, crops show variable genomic signatures of domestication because of differences in their biology and utilization by humans (Meyer *et al* 2012). In particular, domestication affects the level and structure of genetic diversity in crops because selection and genetic drift contributed to strong genetic bottlenecks (Doebley *et al* 2006; Olsen & Wendel, 2013; Sang & Li, 2013; Nabholz *et al* 2014). The geographic expansion of domesticated crops provided the opportunity for gene flow with new crop wild relatives, which further contributed to genetic differentiation from wild ancestors. Such a diversity of phenotypic and genomic changes associated with domestication suggest that the classical model of a single domestication event in a short time span within a small geographic region may not apply to numerous crop plants like barley, apple and olive trees (Besnard & Rubio de Casas, 2015; Cornille *et al* 2012; Poets *et al* 2015). The motivation of the present study was to investigate both the phenotypic and genomic consequences of amaranth cultivation in the light of these concepts.

The genus *Amarantus* L. comprises between 50 and 75 species and is distributed worldwide (Sauer, 1967; Costea & DeMason, 2001). Four species are cultivated as grain amaranths or leaf vegetables (Sauer, 1967; Brenner *et al* 2010). The grain amaranths *Amaranthus caudatus, Amaranthus cruentus* and *Amaranthus hypochondriacus* originated from South and Central America while *A. tricolor* is used as leafy vegetable in Africa. Amaranth is an ancient crop because archaeological evidence in Northern Argentina suggested that wild amaranth seeds were collected and used for human consumption during the initial mid-Holocene (8,000 - 7,000 BP; Arreguez *et al* 2013). In the Aztec empire, amaranth was a highly valued crop and tributes were collected from the farmers that were nearly as high as for maize (Sauer, 1967). Currently, amaranth is promoted as a healthy food because of its favorable composition of essential amino acids and high micronutrient content.

The three grain amaranth species differ in their geographical distribution. *A. cruentus* and *A. hypochondriacus* are most common in Central America, whereas *A. caudatus* is cultivated mainly in South America. In the Andean region, *A. caudatus* grows in close proximity to the two related (i.e., wild) species *A. hybridus* and *A. quitensis*, which are considered as potential ancestors (Sauer, 1967). *A. hybridus* has the widest distribution range from Central to South America while *A. quitensis* is restricted to the central part of South America. *A. quitensis* was tolerated and possibly cultivated in Andean home gardens and used for coloring in historical times.

The history of amaranth cultivation and extent of its domestication are still under discussion (Figure 1). Sauer (1967) proposed two domestication scenarios based on the morphology and geographic distribution of the different species. One scenario postulates three independent domestication events from three different wild ancestors, and another scenario proposes the domestication of *A. cruentus* from *A. hybridus* followed by a migration and intercrossing of *A. cruentus* with *A. powellii* in Central America and an intercrossing of *A. cruentus* with *A. quitensis* resulting in *A. caudatus* in South America. A third scenario was based on genetic markers and suggested that all three cultivated amaranths evolved from *Amaranthus hybridus*, but at multiple locations (Maughan *et al* 2011). Most recently, Kietlinski *et al* (2014) proposed a single domestication of *A. hybridus* in the Andes or in Mesoamerica and a subsequent spatial separation of two lineages leading to *A. caudatus* and *A. hypochondriacus*, or two independent domestication events of *A. hypochondriacus* and *A. caudatus* from a single *A. hybridus* lineage in Central and South America (Figure 1C and D). A more recent analysis based on the phylogeny of the whole Amaranthus genus supports independent domestication of the South American *A. caudatus* and the two Central American grain amaranths from two different, geographically separated lineages of *A. hybridus* as shown in Figure 1D (Stetter & Schmid, 2016).

**Figure 1:**
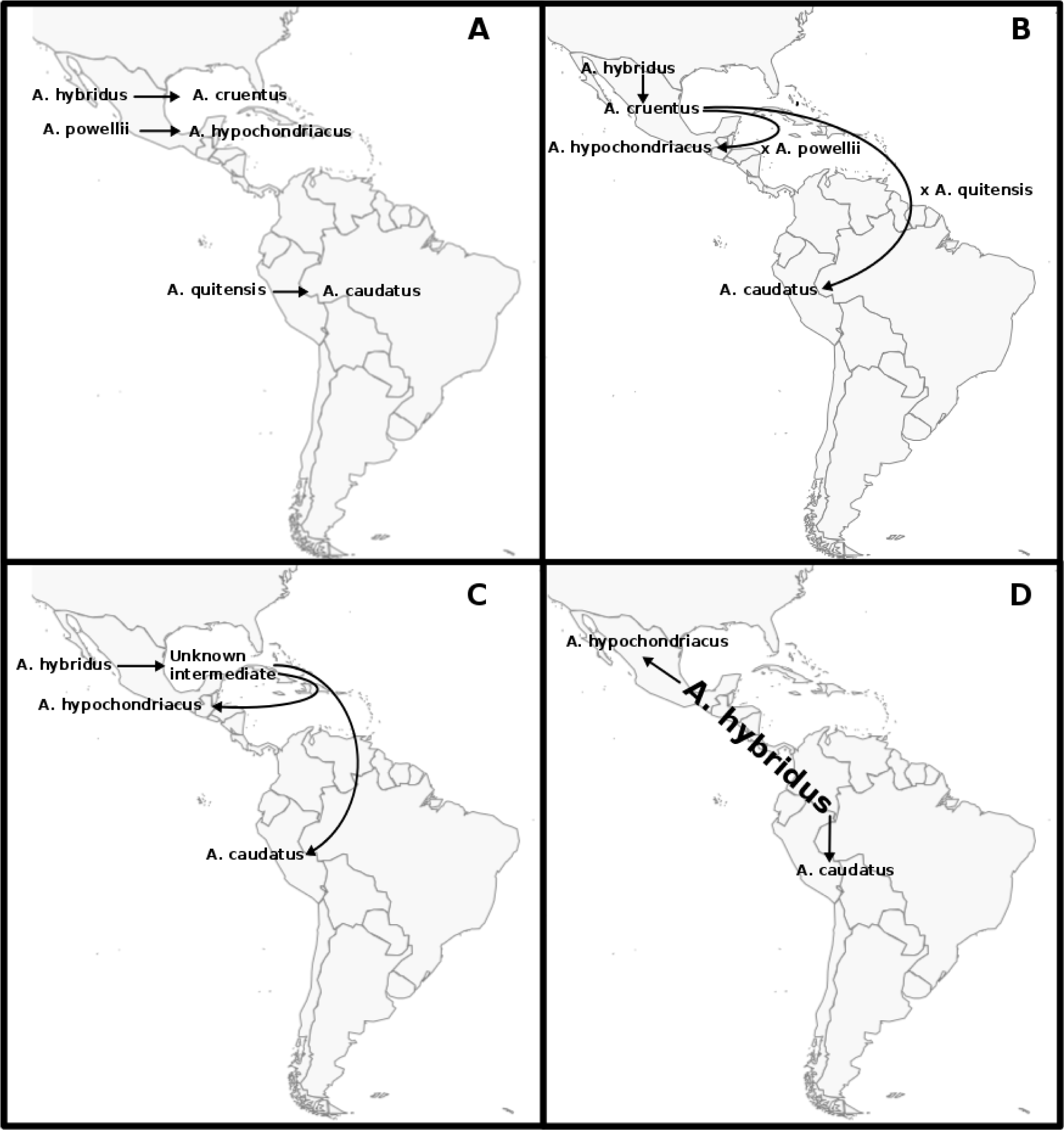
Models of amaranth domestication. (A) Independent domestication of three grain amaranths from different close relatives (Sauer, 1967). (B) Initial domestication from *A. hybridus* and subsequent migration and hybridization with additional close relatives (Sauer, 1967). (C) Single domestication in the Andes or in Mesoamerica and subsequent spatial separation of two lineages leading to *A. caudatus* and *A. hypochondriacus* (Kietlinski *et al* 2014). (D) Two domestication events from a single *A. hybridus* lineage spanning Central and South America (Kietlinski *et al* 2014).

Despite its long history of cultivation and the self-pollinating breeding system, the domestication syndrome of cultivated amaranth is remarkably indistinct because it still shows strong photoperiod sensitivity and has very small shattering seeds (Sauer, 1967; Brenner *et al* 2010). Other crops like maize that were cultivated at a similar time period in the same region exhibit the classical domestication syndrome (Sang & Li, 2013; Lenser & Theißen, 2013). This raises the question whether amaranth is domesticated at all or has a different domestication syndrome, and if the latter is true whether genetic constraints, a lack of genetic variation or (agri-)cultural reasons determined its domestication syndrome. The phenotypic analysis of amaranth domestication is complicated by the taxonomic uncertainty of cultivated amaranth species and their close relatives. Although *A. quitensis* was suggested to be the ancestor of *A. caudatus,* the state of *A. quitensis* as a separate species is under debate. Sauer (1967) classified it as species, but later it was argued that it is the same species as *A. hybridus* (Coons, 1978; Brenner *et al* 2010). However, until today *A. quitensis* is treated as separate species and since genetic evidence for the status of *A. quitensis* as a separate species is based on few studies with limited numbers of markers, this topic is still unresolved (Mallory *et al* 2008; Kietlinski *et al* 2014).

The rapid development of sequencing technologies facilitates the large-scale investigation of the genetic history of crops and their close relatives. Among available methods, reduced representation sequencing approaches such as genotyping-by-sequencing (GBS) allow a genomewide and cost-efficient marker detection compared to whole genome sequencing (Elshire *et al* 2011; Poland *et al* 2012). Despite some biases associated with reduced representation sequencing, GBS and related methods are suitable and powerful approaches for studying interspecific phylogenetic relationships (Cruaud *et al* 2014) and intraspecific patterns of genetic variation in crop plants (Morris *et al* 2013).

We used GBS and genome size measurements to characterize the genetic diversity and relationship of cultivated *A. caudatus* and its possible ancestors *A. quitensis* and *A. hybridus*, and compared patterns of genetic structure with two domestication-related phenotypic traits (seed color and hundred seed weight). For this study, we focussed on the South American amaranth species, because *A. caudatus, A. quitensis* and South American accessions of *A. hybridus* form a clade that is strongly separated from the two Central American grain amaranths in a phylogenetic analysis of the whole genus (Stetter & Schmid, 2016). For this reason, we reasoned that the domestication of *A. caudatus*, which is native to South America, and its relationship to the sympatric relatives, *A. hybridus* and *A. quitensis* can be conducted independently of the Central American amaranth species. Our analysis includes a comparison of genetic diversity and seed-related traits like size and color between cultivated and wild amaranths and analyses the taxonomic relationship and gene flow among species. Our results indicate that *A. caudatus* has a history of domestication that may be considered as incomplete.

## Material and Methods

### Plant material

A total of 119 amaranth accessions of three *Amaranthus* species originating from South America were obtained from the USDA genebank (http://www.ars-grin.gov/npgs/searchgrin.html). Of these accessions, 89 were classified as *A. caudatus*, 17 as *A. hybridus*, seven as *A. quitensis* and six as interspecific hybrids according to the passport information (Table S5). We selected *A. caudatus* accessions based on the altitude of the collection site and focused on high-altitude regions (2,200 to 3,700 m) where amaranth has been cultivated for thousands of years and survived until today since it fell into disuse after the Spanish conquest (Kauffman & Weber, 1990). Therefore, high-altitude accessions may represent a large proportion of the species-wide genetic diversity. The smaller sample sizes of *A. hybridus* and *A. quitensis* accessions reflect that fewer accessions of these species than of *A. caudatus* are available from the USDA genebank. However, the geographic origin of the two wild relatives covers the Andean highlands, which is the distribution range of *A. caudatus*, and we compared the population structure of the sample derived from the genomic data with the passport information to test for consistency between the population structure and geographic origin. Accessions were planted in a field in Nurtingen (Germany), and one young leaf of one representative plant per accession was sampled to avoid the sampling of potential seed cross-contamination. We sampled and sequenced three plants each of 12 accessions independently for quality control.

### Genome size

To compare genome sizes among the three diploid *Amaranthus* species, we measured the genome size of 22 *A. caudatus*, 8 *A. hybridus* and 4 *A. quitensis* accessions. Genome size differences of individuals within species are expected to be low, and we therefore estimated species-specific genome sizes using 25% the total sample of *A. caudatus* and 50% of *A. hybridus* and *A. quitensis* accessions, respectively. Plants were grown for four weeks in the greenhouse before one young leaf was collected for cell extraction. A tomato cultivar *(Solanum lycopersicum* cv Stupicke) was used as internal standard because it has a comparable genome size that has been measured with high accuracy (DNA content = 1.96 pg; Dolezel *et al* 1992). Fresh leaves were cut up with a razor blade and cells were extracted with CyStain PI Absolute P (Partec, Muenster/Germany). Approximately 0.5 cm^2^ of the sample leaf was extracted together with similar area of tomato leaf in 0.5 ml of extraction buffer. The DNA content was determined with CyFlow Space (Partec, Muenster/Germany) flow cytometer and analyzed with FlowMax software (Partec, Muenster/Germany). For each sample, 10,000 particles were measured each time. Two different plants were measured for each accession. The DNA content was calculated as:

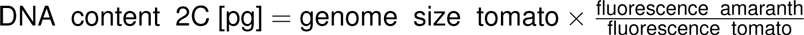

and the genome size (in Mbp) was calculated as:

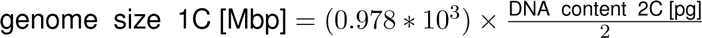

The conversion from pg to bp was calculated with 1pg DNA = 0.978 x 10^9^ bp (Dolezel *et al*, 2003). Means were calculated using R software (Team, 2014) and an ANOVA was performed to infer differences in genome size for the species.

### DNA extraction and library preparation

Genomic DNA was extracted using a modified CTAB protocol (Saghai-Maroof *et al*, 1984). The DNA was dried and dissolved in 50-100 *μ*l TE and diluted to 100 ng/*μ*l for further usage. Two-enzyme GBS libraries were constructed with a modified protocol from the previously described two-enzyme GBS protocol (Poland *et al*, 2012). DNA was digested with a mix of 2 *μ*l DNA, 2 *μ*l NEB Buffer 2 (NEB, Frankfurt/Germany), 1 *μ*l ApeKI (4U/*μ*l, NEB), 1 *μ*l Hindu/ (20U/*μ*l, NEB) and 14 *μ*l ddH_2_O for 2 hours at 37°C before incubating for 2 hours at 75°C. Adapters were ligated with 20 *μ*l of digested DNA 5 *μ*l ligase buffer (NEB), T_4_-DNA ligase (NEB), 4 *μ*l ddH_2_O and 20 *μ*l of adapter mix containing 10*μ*l barcode adapter (0.3 ng/μl) and 10 *μ*l common adapter (0.3ng/*μ*l). Samples were incubated at 22°C for 60 minutes before deactivating ligase at 65°C for 30 minutes. Subsequently, samples were cooled down to 4°C. For each sequencing lane, 5*μ*l of 48 samples with different barcodes were pooled after adapter ligation. Samples of the different species were randomized over the 3 pools and different barcode lengths. The 12 replicated samples were added to each pool. The pooled samples were purified with QIAquick PCR purification kit (Qiagen, Hilden/Germany) and eluted in 50 *μ*l elution buffer before PCR amplification of the pools. The PCR was performed with 10 *μ*l of pooled DNA, 25 *μ*l 2x Taq Master Mix (NEB), 2 *μ*l PCR primer mix (25pmol/*μ*l of each primer) and 13 *μ*l ddH_2_O for 5 min at 72°C and 30 sec at 98°C before 18 cycles of 10 sec at 98°C, 30 sec at 65°C and 30 sec at 72°C after the 18 cycles 5 min of 72°C were applied and samples were cooled down to 4°C. Samples were purified again with QIAquick PCR purification kit (Qiagen) and eluted in 30*μ*l elution buffer. Three lanes with 48 samples per lane were sequenced on an Illumina HighScan SQ with single end and 105 cycles on the same flow cell (see supporting data).

### Data preparation

Raw sequence data were filtered with the following steps. First, reads were divided into separate files according to the different barcodes using Python scripts. Read quality was assessed with fastQC (http://www.bioinformatics.babraham.ac.uk/projects/fastqc/). Due to lower read quality towards the end of the reads, they were trimmed to 90 bp. Low quality reads were excluded if they contained at least one N (undefined base) or if the quality score after trimming was below 20 in more than 10% of the bases. Data from technical replicates were combined and individuals with less than 10,000 reads were excluded from further analysis (Table S5). The 12 replicated samples were used to detect a lane effect with an analysis of variance.

### SNP calling and filtering

Since no high quality reference genome for *Amaranthus* sp. was available for read mapping, we used `Stacks 1.19`, for the *de novo* identification of SNPs in GBS data (Catchen *et al* 2011, 2013). The SNP calling pipeline provided by Stacks `denovounap.pl` was used to call SNPs from the processed data. Highly repetitive GBS reads were removed in the ustacks program with option -t. Additionally, the minimum number of identical raw reads required to create a stack was set to three (`m=3`) and the number of mismatches allowed between loci when processing a single individual was two (`M=2`). Four mismatches were allowed between loci when building the catalog (`n=4`). The catalog is a set of non redundant loci representing all loci in the accessions and used as reference for SNP calling. SNPs were called with the Stacks tool `populations 1.19` without filtering for missing data using option -r 0. One individual, PI 511754, was excluded from further analysis because it appeared to be misclassified. According to its passport information it belonged to *A. hybridus*, but with all clustering methods it was placed into a separate cluster consisting only of this individual, which suggested it belongs to a different species. Therefore, we repeated the SNP calling without this individual. The SNPs were filtered over the whole sample for missing data with vcftools (Danecek *et al* 2011) by allowing a maximum of 60% missing values per SNP position. Given the stringent filtering criteria for SNP calling and the restricted number of *A. quitensis* individuals, we did not filter SNPs by their minor allele frequency for further analysis.

### Inference of genetic diversity and population structure

Nucleotide diversity (n) weighted by coverage was calculated with a Python script that implements the formula of Begun *et al* (2007) which corrects for different sampling depths of SNPs in sequencing data. The confidence interval of π was calculated by bootstrapping the calculation 10,000 times. To account for the difference in sampling between wild and cultivated amaranths, we sub-sampled *A. caudatus* 100 times with the the same number of individuals (23) as used for wild amaranth. The pairwise difference in π between *A. caudatus* and the close relatives was calculated for each site. Mean expected (H_exp_) and observed (H_obs_) heterozygosities based on SNPs were calculated with the R package `adegenet 1.4-2` (Jombart & Ahmed, 2011). The inbreeding coefficient (F) was calculated as:

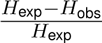

Weir and Cockerham weighted *F*_*ST*_ estimates were calculated with vcftools (Weir & Cocker-ham, 1984; Danecek *et al* 2011). To infer the population structure, we used ADMIXTURE for a model-based clustering (Alexander *et al* 2009) and conducted the analysis with different numbers of predefined populations ranging from *K* = 1 to *K* = 9 to find the value of *K* that was most consistent with the data using a cross-validation procedure described in the ADMIXTURE manual. To avoid convergence effects we ran ADMIXTURE 10 times with different random seeds for each value of K. As a multivariate clustering method, we applied discriminant analysis of principal components (DAPC) implemented in the R-package adegenet (Jombart *et al* 2010; Jombart & Ahmed, 2011) and determined the number of principal components (PCs) used in DAPC with the `optim.a.score` method. To investigate the phylogenetic relationship of the species, we calculated an uncorrected neighbor joining network using the algorithm Neighbor-Net (Bryant & Moulton, 2004) as implemented in the `SpiitsTree4` program (Huson & Bryant, 2006). The Euclidean distance was calculated from the genetic data to construct a neighbor joining tree, which was bootstrapped 1,000 times with the `pegas` R-package (Paradis *et al* 2004). The migration between genetic groups was modeled with TreeMix (Pickrell & Pritchard, 2012). For the TreeMix analysis we used the groups that were identified by ADMIXTURE (K = 5) without an outgroup, and allowed four migration events, as preliminary runs indicates four migration events to be the highest number. The tree was bootstrapped 1,000 times.

### Seed color and hundred seed weight

For each accession we calculated the hundred seed weight (HSW) by weighting three samples of 200 seeds. Seed color was determined from digital images taken with a binocular (at 6.5x magnification) and by visual comparison to the GRIN descriptors for amaranth (http://www.ars-grin.gov/cgi-bin/npgs/html/desclist.pl?i59). There were three colors present in the set of accessions, white, pink, which also indicates a white seed coat and dark brown. To infer how the species, assigned genetic groups or seed color influenced seed size, we conducted an ANOVA. Differences were tested with a LSD test implemented in the R package agricolae (http://tarwi.lamolina.edu.pe/~fmendiburu/).

## Results

### Genome size measurements

Although the genomic history of amaranth species still is largely unknown, genome sizes and chromosome numbers are highly variable within the genus *Amaranthus* (http://data.kew.org/cvaiues/). We therefore tested whether a change in genome size by polyploidization or large-scale insertions or deletions played a role in the speciation history of *A. caudatus* and the two relatives *A. quitensis* and *A. hybridus* by measuring the genome size of multiple individuals from all three species with flow cytometry. The mean genome size of *A. caudatus* was 501.93 Mbp, and the two relatives did not differ significantly from this value (Table 1) indicating that all measured individuals are diploid and that polyploidization did not play a role in the recent evolution of cultivated amaranth.

**Table 1:** Genome size of representative group of individuals for each species. There are no significant differences between genome sizes (p≤0.05). The number of individuals per population is *N* and SD is the standard deviation for each parameter.

| | $N$ | DNA content (pg) | SD | genome size (Mbp) | SD |
| --- | --- | --- | --- | --- | --- |
| *A. caudatus* | 22 | 1.026 | 0.026 | 501.93 | 12.74 |
| *A. hybridus* | 8 | 1.029 | 0.025 | 502.96 | 12.20 |
| *A. quitensis* | 4 | 1.021 | 0.016 | 499.07 | 7.91 |

### SNP identification by GBS

To investigate genome-wide patterns of genetic diversity in *A. caudatus* and its two closest relatives, we genotyped a diverse panel of 119 amaranth accessions from the three species that were initially collected in the Andean region and then obtained from the USDA genebank. The sequencing data generated with a two-enzyme GBS protocol consisted of 210 Mio. raw reads with an average of 1.5 Mio. reads per accession (Supporting information S2). We tested for a lane effect of the Illumina flow cell by sequencing the same 12 individuals on each of the three lanes used for sequencing of the whole sample. An ANOVA of the read number did not show a lane effect (Table S1). Since a high-quality reference genome of an amaranth species was not available, we aligned reads *de novo* within the dataset to unique tags using Stacks (Catchen *et al* 2011). The total length of the aligned reads was 16.6 Mb, which corresponds to approximately 3.3 % of the *A. caudatus* genome. For SNP calling, reads of each individual were mapped to the aligned tags. SNPs were called with parameters described in Materials and Methods, which resulted in 63,956 SNPs and a mean read depth of 40.28 per site. Since GBS data are characterized by a high proportion of missing values, we removed SNPs with more than 60% of missing values. After this filtering step, we obtained 9,485 biallelic SNPs with an average of 35.3 % missing data for subsequent analyses (Figure S1). The folded site frequency spectrum showed an expected distribution but *A. quitensis* had more sites with low frequency due to the restricted number of individuals (Figure S2)

### Inference of population structure

To infer the genetic relationship and population structure of the three amaranth species, we used three different methods that included ADMIXTURE, Discriminant Analysis of Principal Components (DAPC) and phylogenetic reconstruction with an uncorrected neighbor-joining network. The ADMIXTURE analysis with three predefined groups (*K* = 3) that corresponds to the number of *Amaranthus* species included in the study did not cluster accessions by their species, but combined the two relatives *A. hybridus* and *A. quitensis* into a single cluster and grouped the *A. caudatus* accessions into two distinct clusters. Higher values of K did not lead to subdivision of the two close relatives into separate groups that correspond to the species assignment (Figure 2), however, the they were split according to their geographic origin. Crossvalidation showed that K = 5 was most consistent with the data (Figure S3), which produced three different groups of *A. caudatus* accessions that included a few accessions from the close relatives, and two clusters that both consist of *A. hybridus* and *A. quitensis* accessions. These two clusters are not separated by the species assignment but by the geographic origin of accessions because the clusters consist of *A. hybridus* and *A. quitensis* accessions from Peru and Ecuador, respectively, which indicates a strong geographic differentiation among the close relatives.

**Figure 2:**
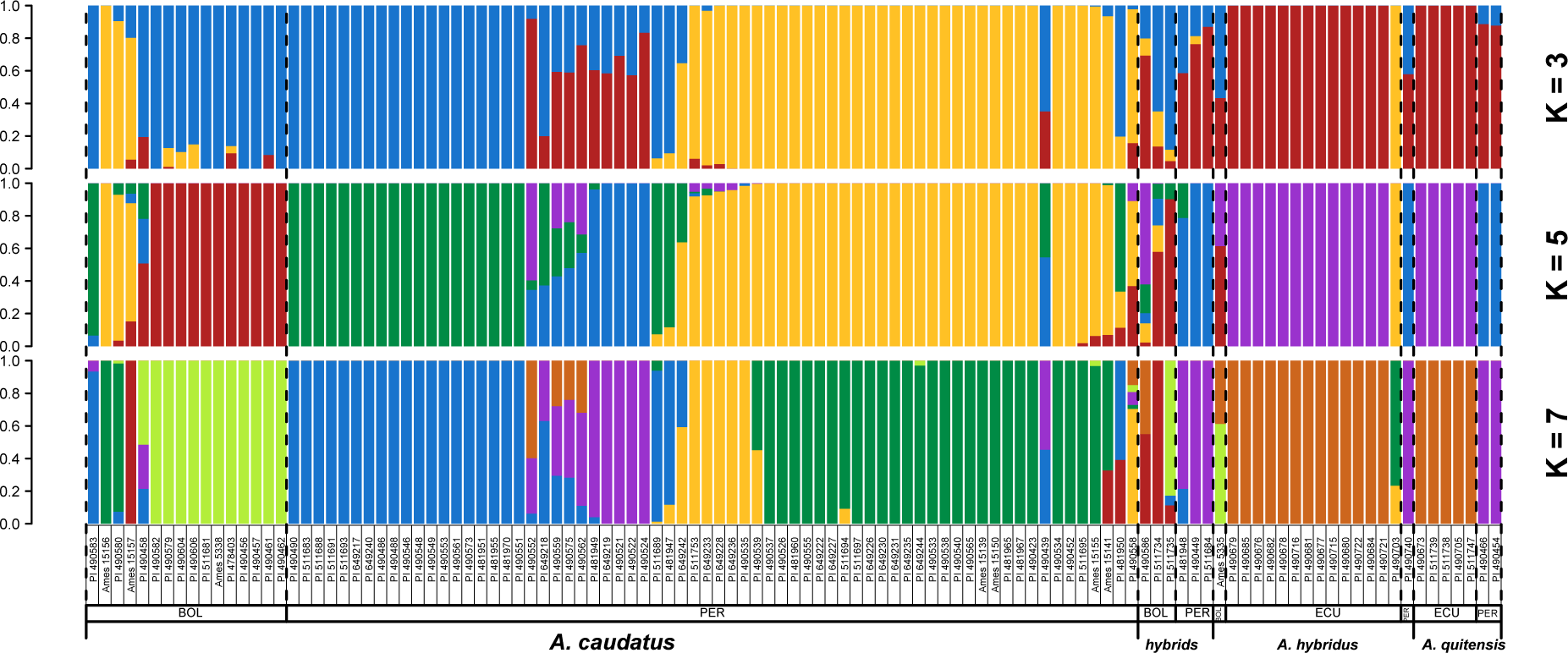
Model based clustering analysis with different numbers of clusters (K=3, 5, 7) with ADMIXTURE. The clusters reflect the number of species in the study (K=3), the number of single populations (species per country of origin, K=7) and the optimal number as determined by cross validation (K=5). Individuals are sorted by species and country of origin (BOL=Bolivia, PER = Peru and ECU = Ecuador) as given by their passport data.

The groups of *A. caudatus* accessions also showed a clear geographic differentiation. The first cluster consisted of individuals from Bolivia (Figures 2 and 3; K = 5, red color). *A. caudatus* accessions from Peru were split into two clusters of which one predominantly represents a region from North Peru (Huari Province; Figures 2 and 3; K = 5, yellow color), whereas the second cluster contains individuals distributed over a wide geographic range that extends from North to South Peru (K = 5, green color). Ten *A. caudatus* accessions from the Cuzco region clustered with the three accessions of the close relatives from Peru (K = 5, blue color). These ten accessions showed admixture with the other cluster of *A. hybridus/A. quitensis* and with a Peruvian cluster of *A. caudatus*. Accessions that were labeled as ‘hybrids’ in their passport data, because they express a set of phenotypic traits of different species, clustered with different groups. ‘Hybrids’ from Bolivia were highly admixed, whereas ‘hybrids’ from Peru clustered with the close relatives from Peru (Figure 2). Taken together, the population structure inference with ADMIXTURE identified a clear separation between the cultivated *A. caudatus* and its to close relatives, and a high level of genetic differentiation among cultivated amaranths with some evidence for gene flow between groups.

**Figure 3:**
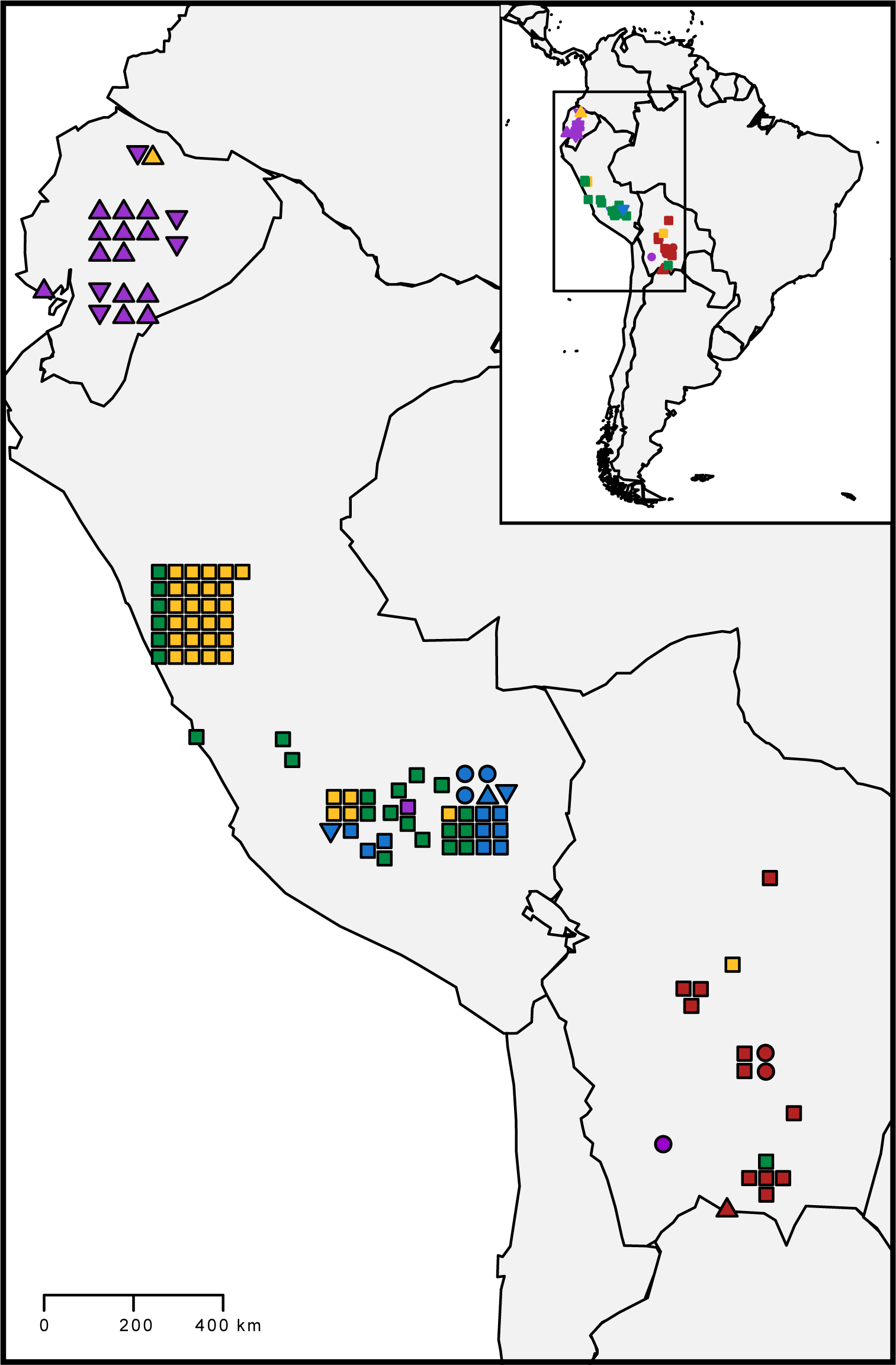
Geographic distribution of accessions for which data was available from passport information. Locations are not exact geographic locations because location data was given as country province. Colors are given by ADMIXTURE with K=5 (Figure 2). Species are indicated by shapes. *A. caudatus* (□), *A. hybridus* (Δ), *A. quitensis* (▽) and hybrids between species (○)

The inference of population structure with a discriminant analysis of principal components (DAPC) and Neighbor-Joining network produced very similar results as ADMIXTURE. The first principal component of the DAPC analysis which we used to cluster accessions based on their species explained 96% of the variation and separated the cultivated *A. caudatus* from its two relatives (Figure S4A). In a second DAPC analysis that was based on information on species and country of origin (Figure S4B) the first principal component explained 55% of the variation and separated most cultivated amaranth accessions from the close relatives. The second principal component explained 35% of the variation and separated the Peruvian from the Bolivian *A. caudatus* accessions.

The phylogenetic network outlines the relationships between the different clusters (Figure 4). It shows two distinct groups of mainly Peruvian *A. caudatus* accessions and a group of accessions with a wide geographic distribution (Figure 3; green color). The latter is more closely related to the Bolivian *A. caudatus* and the close relatives. The strong network structure between these three groups suggests a high proportion of shared ancestral polymorphisms or a high level of recent gene flow. In contrast, *A. caudatus* accessions from Northern Peru are more strongly separated from the other groups (Figure 3; yellow color) and are split into two subgroups, of which the smaller one includes only accessions with dark seeds. In a bifurcating phylogenetic tree, ten cultivated amaranth accessions clustered within the same clade as the close relatives *A. quitensis* and *A. caudatus* (Figure S5). The same clustering was also obtained with ADMIXTURE and *K* = 7 (Figure 2).

**Figure 4:**
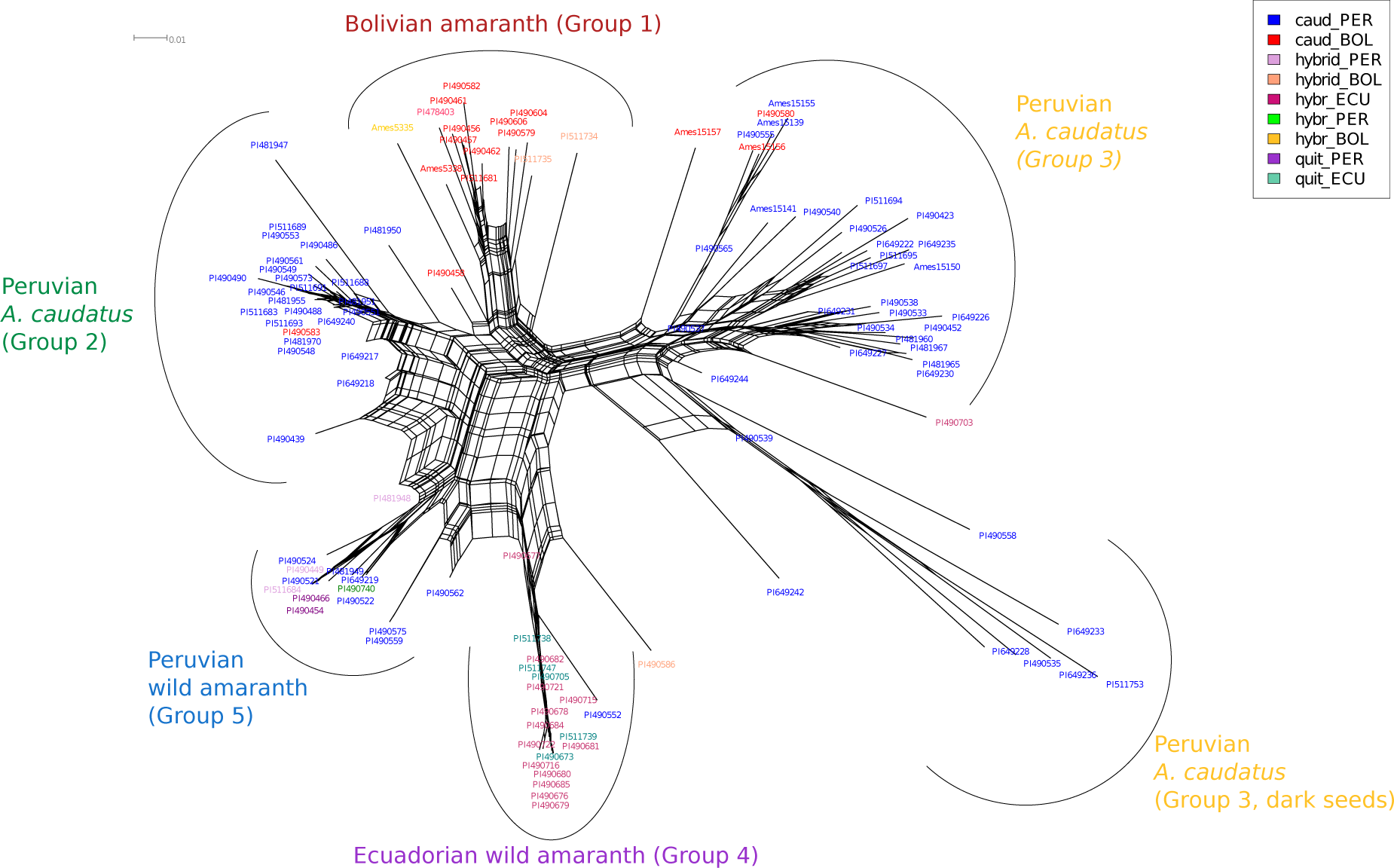
Neighbor-joining network of 113 amaranth accessions from six potential populations. Different colors indicate the species and origin according to gene bank information. *A. caudatus* from Peru (blue) and from Bolivia (red), *A. hybridus* from Ecuador (magenta), from Peru (green) and Bolivia (yellow), *A. quitensis* from Ecuador (turquoise) and Peru (purple) and hybrids between species from Peru (salmon) and Bolivia (light orange). Arches show genetic clusters as inferred with ADMIXTURE (*K* = 5).

To quantify the level of genetic differentiation between the species and groups within *A. caudatus*, we estimated weighted F_*ST*_ values using the method of Weir and Cockerham (Weir & Cockerham, 1984). F_*ST*_ values between *A. caudatus* and *A. hybridus* and *A. quitensis* species were 0.31 and 0.32, respectively (Table 2), and 0.041 between *A. hybridus* and *A. quitensis* based on the taxonomic assignment. The latter reflects the high genetic similarity of the accessions from both species observed above. Within *A. caudatus* subpopulations, the *F*_*ST*_ between *A. caudatus* populations from Peru and Bolivia was 0.132, three times higher than between *A. hybridus* and *A. quitensis*. The above analyses suggested that some individuals may be mis-classified in the passport information, and we therefore calculated F_*ST*_ values of population sets defined by ADMIXTURE. Although such F_*ST*_ values are upward biased, they allow to evaluate the relative level of differentiation between groups defined by their genotypes. The comparison of *F*_*ST*_ values showed that the three *A. caudatus* groups (groups 1-3) are less distant to the group of *A.quitensis/A.hybridus* accessions from Peru (group 5) than from Ecuador (group 4; Table S2). A TreeMix analysis, which is based on allele frequencies within groups (Figure 5), suggests gene flow from the Peruvian *A. caudatus* (group 2) to *A. quitensis* and *A. hybridus* amaranths from Peru (group 5) and, with a lower confidence level, from *A. quitensis* and *A. hybridus* from Ecuador (group 4) into Bolivian *A. caudatus* (group 1), as well as from Bolivian *A. caudatus* to Peruvian *A. caudatus* (Group 2).

**Table 2:** Weir and Cockerham weighted F_*ST*_ estimates between populations based on the taxonomic assignment of their passport data. The group of close relatives are *A. hybridus* and *A. quitensis* taken together.

| | $F_{ST}$ |
| --- | --- |
| *A. caudatus* x *A. hybridus* | 0.319 |
| *A. caudatus* x *A. quitensis* | 0.274 |
| *A. caudatus* x close relatives | 0.322 |
| *A. hybridus* x *A. quitensis* | 0.041 |
| *A. caudatus* (PER) x *A. caudatus* (BOL) | 0.132 |

**Figure 5:**
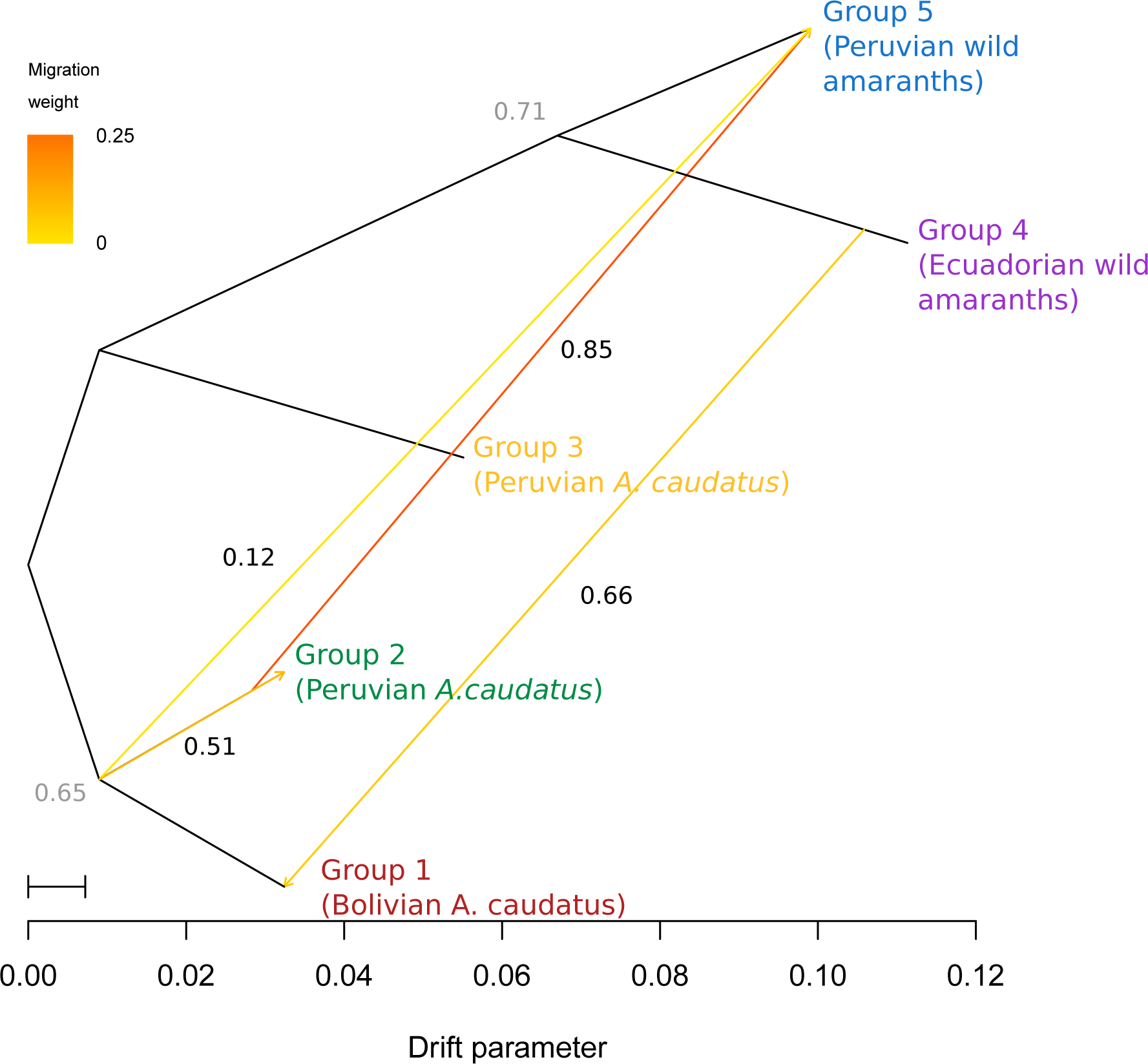
Tree of five genetic clusters of South American amaranths inferred with TreeMix. The genetic clusters which were used to calculate the tree were inferred with ADMIXTURE. Groups 1 to 3 represent *A. caudatus* clusters from Peru and Bolivia, group 4 represents accessions of *A. quitensis* and *A. hybridus* from Ecuador and group 5 wild amaranth from Peru, respectively. The migration events are colored according to their weight. Numbers at branching points and on the migration arrow represent bootstrapping results based on 1,000 runs.

### Analysis of genetic diversity

We further investigated whether domestication reduced genetic diversity in cultivated *A. caudatus* (Table 3). All measures of diversity were higher for *A. caudatus* than its relatives. For example, nucleotide diversity (π) was about two times higher in *A. caudatus* than in the two relatives combined. The diversity values of the accessions classified as ‘hybrids’ showed intermediate values between cultivated amaranth and its relatives supporting their hybrid nature. The inbreeding coefficient, *F*, was highest in the cultivated amaranth but did not differ from the two close relatives if they are combined. In contrast, accessions classified as ‘hybrids’ and *A*. *quitensis* had lower inbreeding coefficients. Within the groups of accessions defined by ADMIXTURE, genetic diversity differed substantially. The close relatives from Ecuador had the lowest (π = 0.00031) while the group from northern Peru showed the highest level of nucleotide diversity (π = 0.00111; Table S3). Figure 6 shows that even though the overall diversity of *A. caudatus* was higher, a substantial proportion of sites were more diverse in the close relatives (*π*_*caud*_ − *π*_*hyb*_/*quit* < 0; Figure 6).

**Figure 6:**
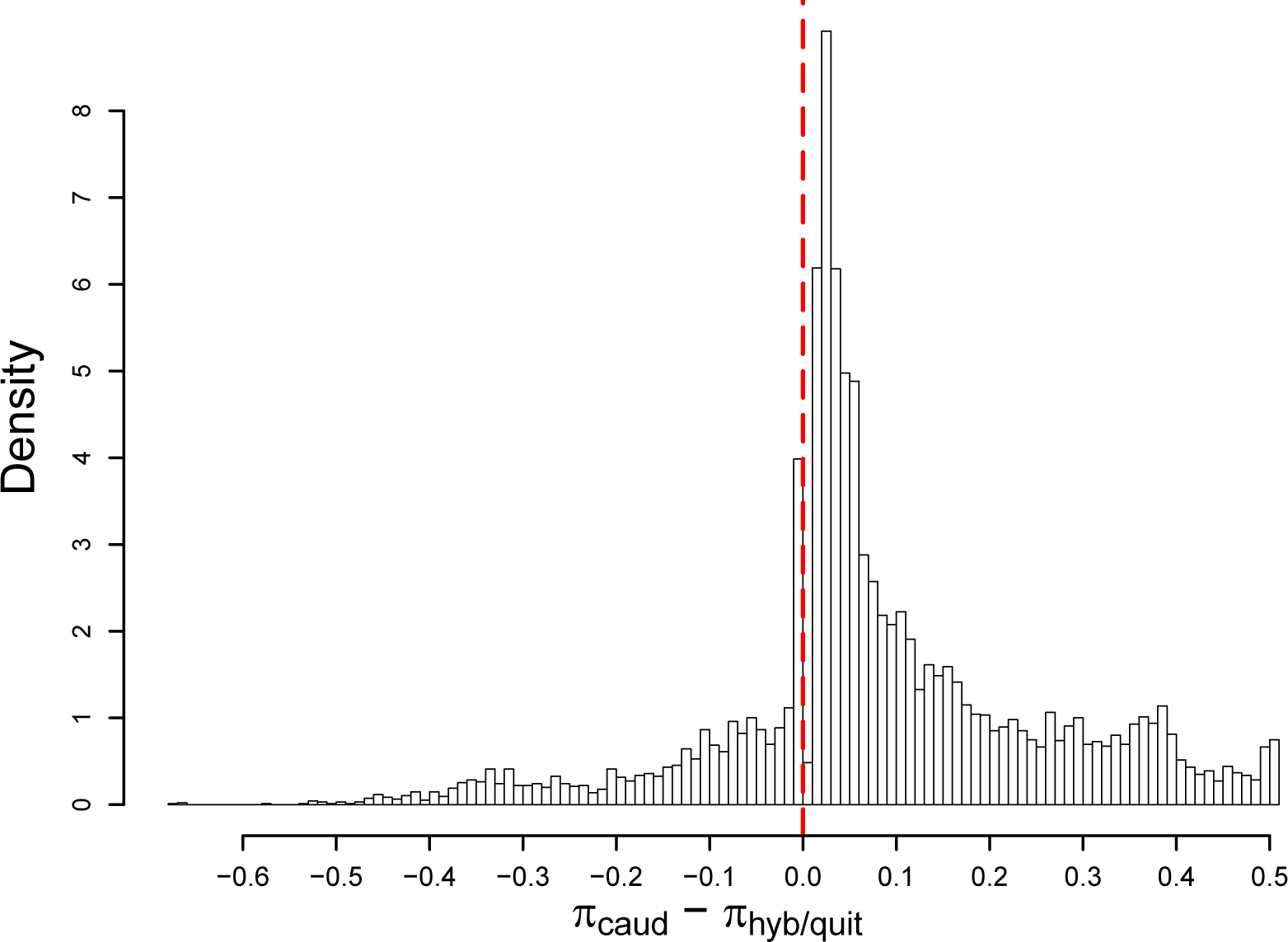
Per site difference in nucleotide diversity (*π*) between cultivated amaranth (*A. caudatus*) and the close relatives *A. hybridus* and *A. quitensis*.

**Table 3:** Genetic diversity parameters for the cultivated *A. caudatus* and the close relatives *A. hybridus* and *A. quitensis. n* is the nucleotide diversity over all sites, CI_n_ is the 95% confidence interval of *π*, *H*_*exp*_ the mean expected heterozygosity for the variant sites and SD_He_ its standard deviation, *H*_*obs*_ the mean observed herterozygosity and SD_*Ho*_ its standard deviation. *F* is the inbreeding coeficient and SD_*F*_ its standard deviation.

| Population | N | $\pi$ | $CI_{\pi}$ | $H_{exp}$ | $SD_{He}$ | $H_{obs}$ | $SD_{Ho}$ | $F$ | $SD_F$ | $\theta_w$ |
| --- | --- | --- | --- | --- | --- | --- | --- | --- | --- | --- |
| *A. caudatus* | 84 | 0.00117 | $\pm 0.00002$ | 0.175 | 0.167 | 0.049 | 0.140 | 0.688 | 0.462 | 0.00123 |
| *A. hybridus* | 16 | 0.00061 | $\pm 0.00001$ | 0.085 | 0.135 | 0.041 | 0.170 | 0.679 | 0.608 | 0.00073 |
| *A. quitensis* | 7 | 0.00059 | $\pm 0.00001$ | 0.076 | 0.169 | 0.040 | 0.170 | 0.451 | 0.763 | 0.00048 |
| Close relatives combined | 23 | 0.00062 | $\pm 0.00002$ | 0.090 | 0.140 | 0.041 | 0.166 | 0.681 | 0.591 | 0.00070 |
| Hybrids | 6 | 0.00091 | $\pm 0.00001$ | 0.112 | 0.179 | 0.060 | 0.173 | 0.436 | 0.645 | 0.00107 |

### Seed color and seed size as potential domestication traits

In grain crops, grain size and seed color are important traits for selection and likely played a central role in domestication of numerous plants (Abbo *et al*, 2014; Hake & Ross-Ibarra, 2015). To investigate whether these two traits are part of the domestication syndrome in grain amaranth, we compared the predominant seed color of the different groups of accessions and measured their seed size. The seeds could be classified into three colors, white, pink and brown. The white and pink types have both a white seed coat, but the latter has red cotyledons that are visible through the translucent seed coat. A substantial number of seed samples (19) from the genebank contained seeds of other color up to a proportion of 20%. One *A. caudatus* accession from Peru (PI 649244) consisted of 65% dark seeds and 35% white seeds in the sample. No accession from the two close relatives *A. hybridus* and *A. quitensis*, or from the hybrid accessions had white seeds, whereas the majority (74%) of *A. caudatus* accessions had white (70%) or pink (4%) seeds, and the remaining (26%) brown seeds (Figure 7 A). We also compared the seed color of groups defined by ADMIXTURE (*K* = 5; Figure 2), which reflect their genetic relationship and may correct for mislabeling of accessions (Figure 7 B). No group had only white seeds, but clusters consisting mainly of *A. hybridus* and *A. quitensis* had no white seeds at all. In contrast to seed color, the hundred seed weight (HSW) of the different *Amaranthus* species did not significantly differ between cultivated *A. caudatus* and the two relatives. The mean HSW of *A. caudatus* was 0.056 g and slightly higher than the HWS of *A. hybridus* (0.051 g) and *A. quitensis* (0.050 g; Figure 7 C and Table S4). Among the groups identified by ADMIXTURE (*K* = 5), one group showed a significantly higher HSW than the other groups, while the other four groups did not differ in their seed size. The group with the higher HSW consisted mainly of Bolivian *A. caudatus* accessions and had a 21 % and 35 % larger HSW than the two groups consisting mainly of Peruvian *A. caudatus* accessions, respectively (Figure 7 D). An ANOVA also revealed that seed color has an effect on seed size because white seeds are larger than dark seeds (Table 4).

**Figure 7:**
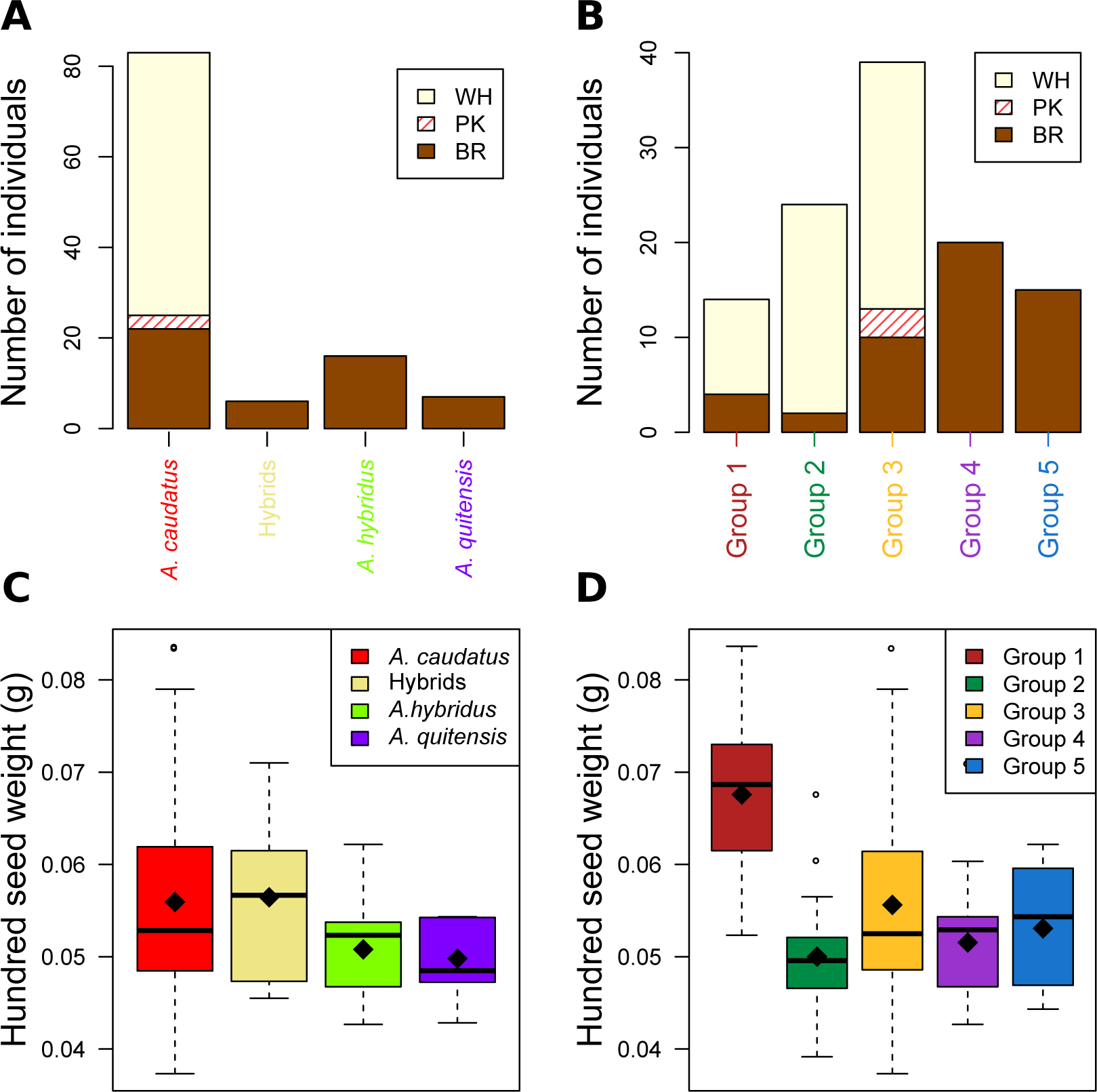
Predominant seed color (**A,B**) and hundred seed weight (**C,D**) by *Amaranthus* species (**A,C**) and groups identified with ADMIXTURE for K = 5 (B,D) where group 1 (red) resembles *A. caudatus* from Bolivia, group 2 (green) and 3 (yellow) *A. caudatus* from Peru, group 4 (purple) represents the close relatives *A. quitensis* and *A. hybridus* from Ecuador and group 5 (blue) from Peru, respectively. Seed colors were white (WH), pink (PK) and dark brown (BR). While there were no significant differences in seed size between the species, group 1 had significantly higher hundred seed weight (p ≤ 0.05) than the other groups.

**Table 4:**
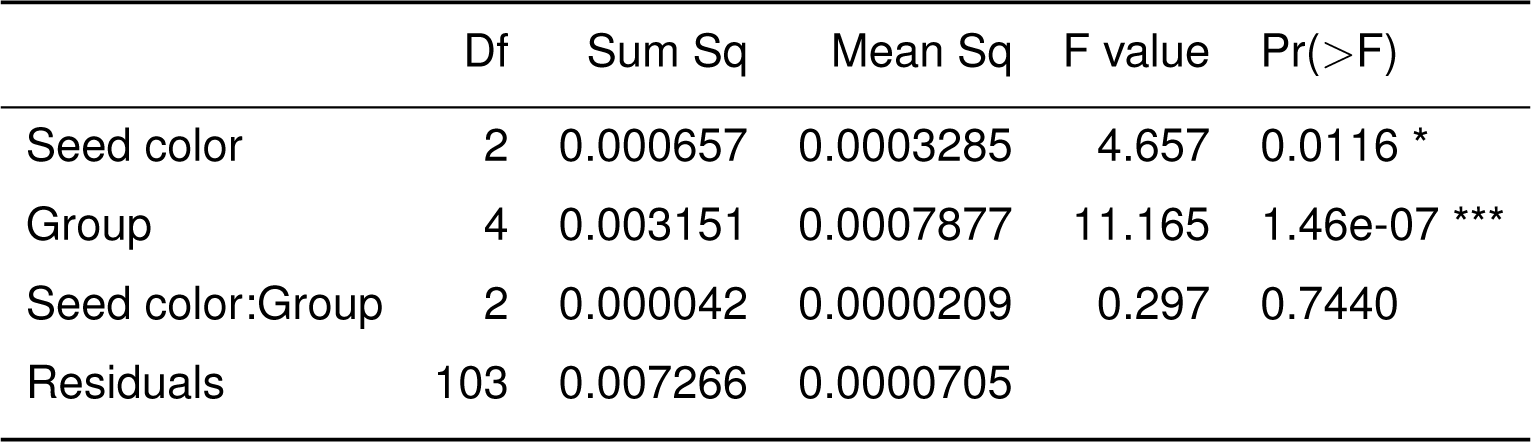
Analysis of variance for the hundred seed weight in dependence of the seed color and population as determined by ADMIXTURE

|  | Df | Sum Sq | Mean Sq | F value | Pr(>F) |
| --- | --- | --- | --- | --- | --- |
| Seed color | 2 | 0.000657 | 0.0003285 | 4.657 | 0.0116 * |
| Group | 4 | 0.003151 | 0.0007877 | 11.165 | 1.46e-07 *** |
| Seed color:Group | 2 | 0.000042 | 0.0000209 | 0.297 | 0.7440 |
| Residuals | 103 | 0.007266 | 0.0000705 |  |  |

## Discussion

### Genotyping-by-sequencing of amaranth species

The genotyping of cultivated amaranth *A. caudatus* and two close relatives *A. quitensis* and *A. hybridus* revealed a strong genetic differentiation between both groups and a high level of genetic differentiation within cultivated *A. caudatus*. We based our sequence assembly and SNP calling on a *de novo* assembly of GBS data with Stacks because currently no high quality reference sequence of these species is available. Stacks allows SNP calling without a reference genome by constructing a reference catalog from the data and includes all reads in the analysis (Catchen *et al* 2011). *De novo* assembled fragments without a reference are unsorted and can not be used to investigate genetic differentiation along along the genomic regions, but they are suitable for analysing genetic diversity and population structure (Catchen *et al* 2013). GBS produces a large number of SNPs (Poland *et al* 2012; Huang *et al* 2014), albeit with a substantial proportion of missing values. Missing data lead to biased estimators of population parameters such as n and *d*_*w*_ (Arnold *et al* 2013) and need to be accounted for if different studies are compared. Additionally, variable error rates in different GBS data sets can inflate differentiation estimates (Mastretta-Yanes *et al* 2015). The comparison of accessions and groups within a study is possible, however, if all individuals were treated with the same experimental protocol. We filtered out sites with high levels of missing values to obtain a robust dataset for subsequent population genomic analysis. The SNPs were called based on the total sample without accounting for the species which should not bias diversity estimates. Since a smaller sample from the close relatives may underestimate their diversity compared to cultivated *A. caudatus*, we compared diversity estimates by repeated random sampling of 23 out of 84 *A. caudatus* accessions and calculating n from the smaller sample. Diversity estimates of the smaller *A. caudatus* did not differ from the full sample and estimates were in all cases higher than in the close relatives (Figure S6). We conclude that the different sample sizes of the two groups do not introduce a bias on diversity estimates.

### Genetic relationship of *A. quitensis* and *A. hybridus*

Coons (1978) suggested that *A. quitensis* is the same species as *A. hybridus*, but in the genebank passport data *A. quitensis* is still considered as a separate species. The taxonomic differentiation between the two species rests on two minor morphological trait, namely the shape of the tepals and the short utricles, which are very small and prone to misidentification (Sauer, 1967; Adhikary & Pratt, 2015). The high phenotypic similarity of *A. quitensis* and *A. hybridus* is supported by the GBS data because accessions from the two species are closely related. They are not separated by their species assigment but cluster into two groups that both consist of accessions from the two species and reflect their geographic origin from Peru and Ecuador, respectively. The F_*ST*_ value between *A. quitensis* and *A. hybridus* was lower than between the two *A. caudatus* groups from Peru and Bolivia (Tables 2 and S2). The highly similar genome sizes of all three diploid species is consistent with genetic relationship inferred from the GBS data and indicates that large-scale genomic changes like polyploidization events did not occur in the recent history of these species. For comparison, other species in the genus *Amaranthus* have very different genome sizes due to variation in chromosome numbers and ploidy levels (Baohua & Xuejie, 2002; Rayburn *et al* 2005).

In contrast to our analysis, Kietlinski *et al* (2014) found stronger evidence for a genetic differentiation between *A. hybridus* and *A. quitensis* based on the 11 SSR markers. However, their data also show that both species are distinct groups that do not cluster their species assignment but by geographic origin. These differences may result from the different marker types (SNPs vs. SSRs) and a different sample composition because our sample consists of accessions from the Andean region, whereas Kietlinski *et al*. included putative wild amaranth accessions with little geographic overlap between the two species. The groups of *A. hybridus* and *A. quitensis* accessions from Peru and Ecuador show a high level of differentiation (F_*ST*_ = 0.579; Table S2), which is similar to the differentiation between one of two Peruvian *A. caudatus* groups and the *A. hybridus/A.quitensis* accessions from Peru (*F*_*ST*_ = 0.553). Although the sample size of *A. quitensis* and *A. hybridus* is small, genetic differentiation between species should be stronger than between individuals within species in the ADMIXTURE and phylogenetic analyses. In summary, our analysis and the work of Kietlinski *et al* (2014) show that *A. quitensis* and *A. hybridus* do not have a simple genetic relationship that follows species assignment. The high level of intraspecific differentiation in both cultivated amaranth and their relatives is relevant for investigating domestication because the genetic distance between groups of cultivated amaranth is related to the geographic distance of the putative wild ancestors. Therefore, future studies of these two close relatives of the grain amaranths should include large number of accessions from the whole species range to model genetic differentiation within the two species as well as the relationship between species.

### Diversity of South American amaranth

In numerous crops, domestication was associated with a decrease in genome-wide levels of diversity due to bottleneck effects and strong artificial selection of domestication traits (Gepts, 2014). Under the assumption that the cultivated grain amaranth *A. caudatus* was domesticated, genetic diversity should be reduced compared to the two close relatives. In contrast, the overall genetic diversity in our sample of cultivated amaranths was higher than in the two close relatives. The distribution of diversity between the GBS fragments includes genomic regions with reduced diversity in *A. caudatus*, which may reflect selection in some genomic regions (Figure 6). Without a reference genome it is not possible to position reads on a map to identify genomic regions that harbor putative targets of selection based on an inference of the demographic history. Despite the indirect phenotypic evidence for selection, the higher genetic diversity of cultivated grain amaranth may result from a strong gene flow between cultivated amaranths and its relatives. Gene flow between different amaranth species has been observed before (Trucco *et al* 2005) and is also consistent with the observation of six highly admixed accessions classified as ‘hybrids’ in the passport data and which appear to be interspecific hybrids (Figure 2 and Table 3). Gene flow between *A. caudatus* and the relatives *A. quitensis* and *A. hybridus* in different areas of the distribution range, not only from populations included in this study, could explain a higher genetic diversity in cultivated amaranth. This is also consistent with the strong network structure (Figure 4) and the TreeMix analysis (Figure 5). In summary, cultivated *A. caudatus* is unusual in its higher overall genetic diversity compared to populations of its putative wild ancestors originating from the same geographic region. The high genetic diversity of *A. caudatus* is in contrast to other domesticated plants and suggests that a domestication bottleneck in its cultivation history absent (i.e., no domestication), very weak or masked by recurrent gene flow. We consider these results to be robust, because in comparison to previous work (Maughan *et al* 2009, 2011; Khaing *et al* 2013; Jimenez *et al* 2013; Kietlinski *et al* 2014), our study includes a larger number of accessions and more genetic markers. This allowed us to assess the genetic diversity and population structure of South American amaranth on a genome-wide basis, but we note that a more complete geographic sampling of all cultivated amaranths and their relatives is required for a complete understanding of amaranth history.

### Evidence for a weak domestication syndrome

Despite their long history of cultivation, diverse uses for food and feed and a high importance for the agriculture of ancient cultures (i.e., *A. hypochondriacus* during the Aztec period), grain amaranths do not display the classical domestication syndrome as strongly as other crops (Sauer, 1967). Cultivated amaranth shows morphological differentiation from putative wild ancestors like larger and more compact inflorescences (Sauer, 1967) and a level of genetic differentiation (Table 2) which is comparable to other domesticated crops and their wild relatives (Sunflower: F_*ST*_=0.22 (Mandel *et al* 2011); common bean: 0.1-0.4 (Papa *et al*, 2005), pigeonpea: 0.57-0.82 (Kassa *et al* 2012)). However, the numerous amaranth flowers mature asynchronously and produce very small seeds that are shattered (Brenner *et al* 2010). All putative wild amaranths have dark brown seeds, whereas the predominant seed color of cultivated grain amaranth is white, which suggests that selection for seed color played a role in the history of the latter. Dark-seeded accessions are present in all three groups of *A. caudatus* defined by the genotypic data, which indicates that white seed color is not a fixed trait. Seed sizes between cultivated amaranth and its relatives are not significantly different with the exception of white-seeded *A. caudatus* accessions from Bolivia (Figure 7), which have larger seeds. The larger seeds in this group and of white seeds in general (Table 4) indicates past selection for domestication-related traits, but only in specific geographic regions or in certain types of amaranth, and not in the whole cultivated crop species.

These findings suggest that some selection occurred in the history of amaranth cultivation that may reflect domestication. Possible explanations for the incomplete fixation of domestication traits in South American grain amaranth include weak selection, genetic constraints or ongoing gene flow. First, weak selection of putative domestication traits indicate that they were not essential for cultivation. Although white seeds are predominant in cultivated amaranth and most likely a selected trait, other seed colors may have been preferred for different uses. Since amaranths are also an important leaf vegetable in Mexico, the grain use of *A. caudatus* may not have been a main target of selection during domestication, thereby allowing a diversity of seed traits due to weak or incomplete selection. Second, genetic constraints may limit phenotypic variation in domestication traits. In contrast to genes with strong pleiotropic effects or epistatic interactions, domestication genes that are part of simple molecular pathways, have minimal pleiotropic effects, and show standing functional genetic variation have a higher chance of fixation by selection (Doebley *et al* 2006; Lenser & Theißen, 2013). Numerous genes with these characteristics were cloned and characterized in major crops like rice, barley and maize. They contribute to the distinct domestication syndrome such as a loss of seed shattering, larger seed size and compact plant architecture. The molecular genetics of amaranth domestication traits remains unknown, but the absence a strong domestication syndrome may reflect genetic constraints despite a long period of cultivation. A third explanation is ongoing gene flow between cultivated amaranth and its relatives that may prevent or delay the formation of a distinct domestication syndrome and contributes to the high genetic diversity (Table 3), similar seed size (Figure 7 C), and the presence of dark seeds (Figure 7) in cultivated amaranth. Both historical and ongoing gene flow are likely because amaranth has an outcrossing rate between 5% and 30% (Jain *et al* 1982; Stetter *et al* 2016). In South America, cultivated amaranth and its relatives are sympatric over wide areas and the latter were tolerated in the fields and home gardens with *A. caudatus* (Sauer, 1967), where they may have intercrossed. Gene flow between wild and domesticated plants has also been observed in maize and teosinte in the Mexican highlands, but did not have a major influence on the maize domestication syndrome (Hufford *et al* 2013). Further support for ongoing gene flow in amaranth is given by the presence of hybrids and admixed accessions in our sample with evidence for genetic admixture and dark seeds that demonstrate the phenotypic effects of introgression. Since the dark seed color is dominant over white color (Kulakow *et al* 1985) and *A. caudatus* is predominantly self-pollinating, dark seeds could have efficiently removed by selection despite gene flow. Additionally, amaranth was grown in small plots in the Andean highlands, which favors fixation of traits (Kauffman & Weber, 1990). Thus, gene flow is a plausible explanation for the absence of a distinct domesti-cation syndrome.

Although our sample does not include *A. hypochondriacus* or *A. cruentus* accessions, our data are consistent with the model by Kietlinski *et al* (2014) who proposed the domestication of *A. caudatus* and *A. hypochondriacus* from different *A. hybridus* lineages in Central and South America (Figure 1D). Gene flow between *A. caudatus* and its close relative *A. quitensis* in the Southern distribution range (Peru and Bolivia) may explain the higher genetic diversity of the latter despite a strong genetic differentiation.

## Conclusions

The genotypic and phenotypic analysis of cultivated South American grain amaranth and its close relatives suggests that *A. caudatus* is an incompletely domesticated crop species. Key domestication traits such as the shape of inflorescences, seed shattering and seed size are rather similar between cultivated amaranths and their close relatives and there is strong evidence of ongoing gene flow between these species despite selection for domestication-related traits like white seeds. Grain amaranth is an ancient crop of the Americas but genomic and phenotypic signatures of domestication differ from other, highly domesticated crops that originated from single domestication events like maize (Hake & Ross-Ibarra, 2015). In contrast, the history of cultivated amaranth may include multiregional, multiple and incomplete domestication events with frequent and ongoing gene flow from sympatric relatives, which is more similar to the history of species like rice, apple or barley (Londo *et al* 2006; Cornille *et al* 2012; Poets *et al* 2015). The classical model of a single domestication in a well-defined center of domestication may not sufficiently reflect the history of numerous ancient crops. Our study further highlights the importance of a comprehensive sampling to study the domestication of amaranth. The three cultivated amaranths and all close relatives should be included in further studies for a full understanding of amaranth domestication and its broader implications for crop plant domestication.

## Acknowledgments

We thank David Brenner (USDA-ARS) and Julie Jacquemin for discussions and Elisabeth Kokai-Kota for support with the GBS library preparation and sequencing. The work was funded by an endowment of the Stifterverband fur die Deutsche Wissenschaft to K. J. S.

## Data Accessibility

The original genomic data are available on the European Nucleic Archive (ENA) under the study accession number PRJEB13013. Scripts and phenotypic raw data are available under http://dx.doi.org/10.5061/dryad.m5kk3 on Dryad (http://datadryad.org/).

## Author Contributions

M.G.S. and K.J.S. designed research; M.G.S. and K.J.S. performed research; T.M. contributed analytic tools; M.G.S. analyzed data; and M.G.S. and K.J.S. wrote the paper.

